# Miniaturized widefield microscope for high speed in vivo voltage imaging

**DOI:** 10.1101/2025.08.04.668551

**Authors:** Catherine A. Saladrigas, Forest Speed, Alec Teel, Mo Zohrabi, Eduardo J. Miscles, Gregory L. Futia, Larry V. Baker, Ye Zhang, Ioannis Kymissis, Victor M. Bright, Cristin G. Welle, Diego Restrepo, Juliet T. Gopinath, Emily A. Gibson

## Abstract

Functional imaging in freely moving animals with genetically encoded voltage indicators (GEVIs) will open new capabilities for neuroscientists to study the behavioral relevance of neural activity with high spatial and temporal precision. However, miniaturization of an imaging system with sufficient collection efficiency to resolve the small changes in fluorescence yield from voltage spikes, as well as development of efficient image sensors that are sufficiently fast to capture them, has proven challenging. We present a miniaturized microscope designed for voltage imaging, with a numerical aperture of 0.6, 250 μm field of view and 1.3 mm working distance that weighs 16.4 g. We show it is capable of imaging in vivo voltage spikes from Voltron2 with a spike peak-to-noise ratio >3 at a framerate of 530 Hz.

## 1. Introduction

High-speed imaging of neural activity using genetically encoded voltage indicators is an exciting tool in neuroscience that enables precise measurement of firing rates and synchronized activity. Miniature microscopes have been developed to correlate activity across neural circuits with behavior in freely moving animal models, although they have been used predominately for imaging calcium indicators [1–8]. Imaging with genetically encoded calcium indicators (GECIs) is currently the standard for most functional fluorescence imaging applications [9–14]. However, slow calcium efflux kinetics degrade the temporal resolution at which neuronal signals can be monitored using GECIs and require post-processing for the detection of underlying spike trains involved in neural computation that may not be accurate [9]. Genetically encoded voltage indicators (GEVIs) have risen as a new tool to enable neuroscience researchers to study spiking activity related to neural computation in awake rodents [15–19]. While both GECIs and GEVIs produce a time-dependent fluorescent signal that correlates to neural activity, GEVIs are directly sensitive to membrane voltage changes and therefore provide a more accurate temporal response with millisecond rise and fall times than measuring calcium changes [20]. However, voltage imaging in freely moving animals remains limited due to high frame rate (>400 Hz) and collection efficiency requirements for accurate detection of action potentials [21]. For researchers to attain the most accurate understanding of neural computation, novel optoelectronic technologies and computational methods must be developed that enable experiments with GEVIs.

To date, a variety of miniaturized microscopes (miniscopes) have been developed to perform *in vivo* studies on freely moving animals [22–29]. Miniature microscopes are an innovation in which microscopy techniques are implemented in a small, lightweight form factor so that they can mounted on the heads of animals like mice. Without any counterweighting, mice can support devices up to ∼4 grams, while rats can carry up to 35 gram devices on their heads [30, 31]. In particular, widefield miniscopes have been widely used for imaging with GECIs [1–8]. Widefield imaging with Voltron2 requires lower excitation intensities by 2-3 orders of magnitude than techniques like two-photon microscopy to reach the same signal-to-noise ratio (SNR), does not require an onboard scanning element, and frame rate is predominantly limited by camera speeds which can now reach several kHz [32, 33]. Furthermore, the Voltron2 voltage indicator is better suited for one-photon excitation, and thus for widefield miniscopes, over two-photon designs due to the relatively high intensities (an order of magnitude higher than calcium indicators) required to produce enough fluorescence to resolve the voltage spikes [32, 33]. A notable example of a widefield miniscope is the open source UCLA Miniscope and its derivatives, which have been successful in a variety of calcium imaging experiments thanks to a light weight (< 4 grams), and customizability [34]. However, these devices are not optimized for high numerical aperture as they are designed for air and cannot operate at high framerates from a combination of lower excitation intensity from an LED source and cannot acquire images at 100’s of Hz. Experiments that require freely behaving rodents remain the most difficult to adapt to voltage imaging.

Challenges with extending voltage imaging to miniature microscopes arise from stringent size and weight requirements for freely moving animal studies coupled with demanding sensor specifications and collection efficiency. For accurate detection of underlying voltage spikes, one must distinguish a slight change in baseline fluorescence emitted from in-focus cells above fluctuations caused by contamination from out-of-focus background and stochastic noise. While recent voltage indicators such as Voltron2 enable spike ΔF / F of ∼9% for in-focus cells (as governed by the efficiency of the indicator), baseline saturation caused by out-of-focus fluorescence may further reduce spike ΔF / F to only 1-3% [15]. This necessitates high collection efficiency, contrast between in- and out-of-focus fluorescence, and a camera with sufficient sensitivity, pixel size and framerate to provide spatiotemporal resolution for detection of individual action potentials with cellular resolution. These points all prove to be at odds with miniaturization. In general, collection efficiency and depth of field improve with increasing numerical aperture (NA) of an imaging system. NA is increased with larger optical components. However, this intrinsically increases weight, which we seek to minimize for freely moving animal experiments. Furthermore, there are limited camera options with the appropriate framerate and pixel size in a lightweight form factor.

Here, we present the first miniaturized form factor widefield microscope, MiniVolt, designed for voltage imaging. Our device is optimized for a numerical aperture > 0.5 to improve fluorescence collection efficiency with the Sony IMX568 image sensor with sufficient quantum efficiency (80% at 550 nm), pixel size (2.2 × 2.2 microns) and framerate (> 500 Hz) to resolve voltage spikes. MiniVolt is constructed entirely from commercially available components including optics and the camera. To demonstrate the capabilities of our miniscope, we perform imaging in NDNF-Cre transgenic mice expressing Voltron2 in the visual cortex, with framerates up to 530 Hz. On average we have SNR and spike peak-to-noise ratio (PNR) comparable to voltage imaging experiments with benchtop optoelectronics, where our PNR is consistently >3.

## 2. Methods

### 2.1. MiniVolt Imaging System

All experiments presented use a 532 nm laser (Coherent, Verdis V5) for the fluorescence excitation source. Before coupling into a multimode fiber (custom Thorlabs patch cable, 800 μm core, 0.39 NA), the light is passed through a series of N-BK7 static diffusers (Thorlabs, DG10-220 and DG10-1500) to reduce speckle from laser coherence. The multimode fiber is press fit into the miniature microscope. The miniature device is comprised of four commercial lenses from Edmund optics, a dichroic (Chroma T550lpxr) to separate excitation and emission, and an emission filter (Chroma ET570lp) before the sensor. Lens L_1_ (see Fig. 1a and b) operates approximaty as a tube lens, collimating the excitation fiber output and focusing the emission onto the sensor, with lenses L_2_ and L_3_ acting together as an objective module. The dichroic and emission filter are chosen to maximize collection efficiency for Voltron2_552_. In addition to using slightly larger lenses (6-9 mm) than are often used in devices such as the UCLA miniscope [34] or the SIMscope3D (∼ 3 mm) [23], our device is designed to work as a water immersion microscope, enabling a 0.6 NA. This is the highest NA to date in a widefield miniscope. Other specifications of our imaging design include a 250 micron diameter field of view (FOV), a working distance of 1.5-1.6 mm, and a 2.5× magnification. The FOV is determined by imaging a fluorescent grid (Max Levy, DA113), as seen in Fig. 1c, which also shows a uniform illumination field. Furthermore, we compare the contrast of the grid target on the benchtop microscope and MiniVolt and find the respective contrasts are 32.3% and 41.2%. The simulated point spread function is presented in Fig. S1.

**Fig. 1.**
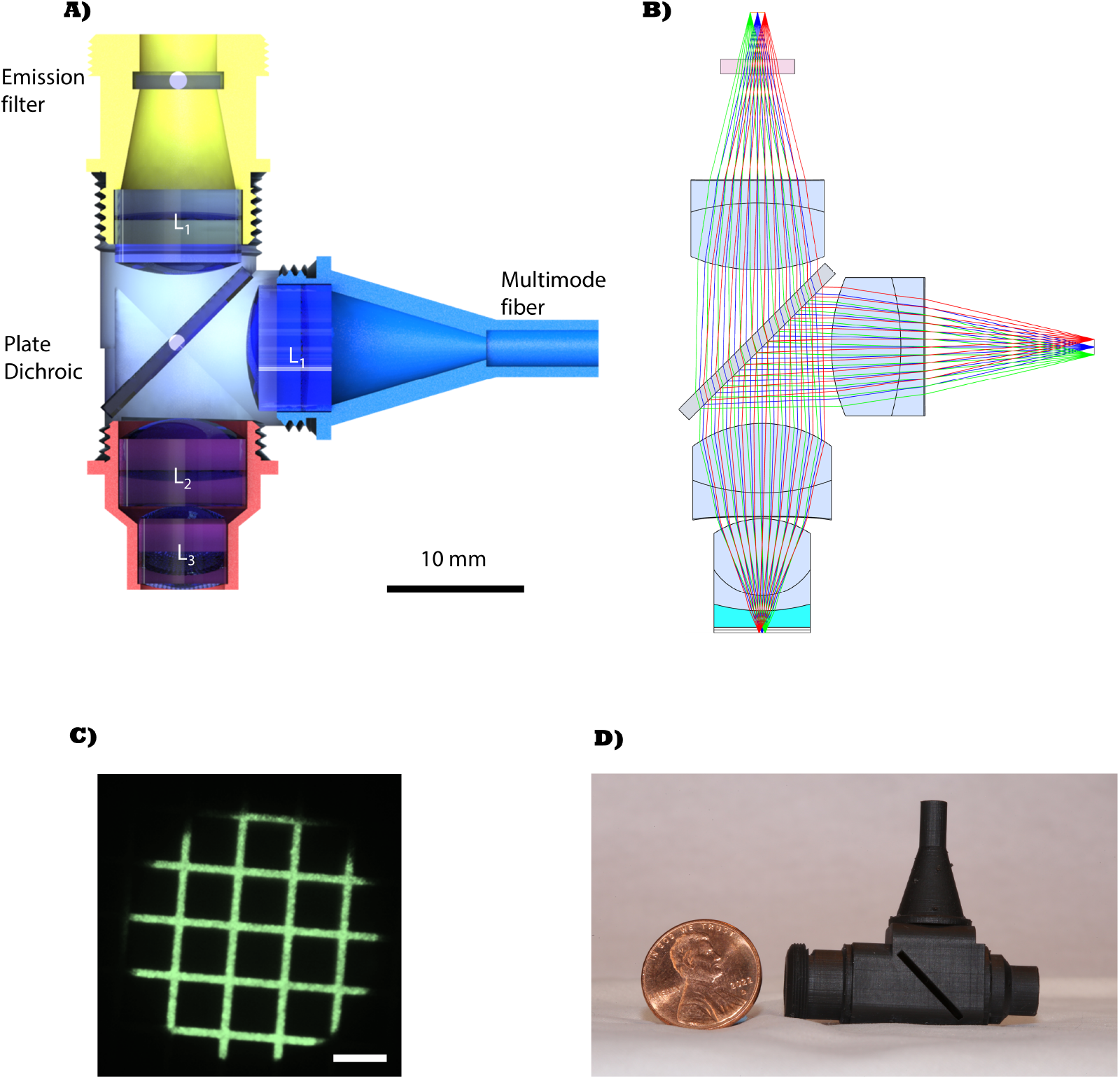
Overview of MiniVolt. (a) CAD schematic showing a cross-sectional view of MiniVolt. (b) Zemax optical design for the emission path of MiniVolt. (c) Image of fluorescent grid target taken with MiniVolt. Scale bar corresponds to 50 microns. (d) Photo of MiniVolt next to a penny for size reference. Optics in (a) and (b) correspond to following: L_1_-Edmund Optics 49-657, L_2_-Edmund Optics 49-656, L_3_-Edmund Optics 49-924, Emission filter-Chroma ET570lp (6×6×1 mm), and Dichroic-Chroma T550lpxr (14×14×1 mm)

The device is enclosed in a custom 3D printed housing. The housing has a modular design with each arm threading together on a common body, as seen in the cross section in Fig. 1a. A picture of the device is shown in Fig. 1d next to a US penny for size reference. This results in a robust housing design with the ability to finetune the separation between each optical element. Finally, the housing threads into the camera (Ximea MU051MG-SY). Our total device weight is 16.4 grams, comprised of 4.6 grams from the housing, 4.9 grams from the optics, and 6.9 grams from the camera. In Table 1, we compare specifications of MiniVolt to other widefield miniscope designs. Although our device weighs more than other designs targeted for mice, it is unmatched in numerical aperture (and therefore collection efficiency) and camera frame rate. Furthermore, our weight would be suitably low (<35 grams) for freely moving rats.

**Table 1.** Comparison of miniature microscope specs

|  | Weight (g) | NA | FOV | Working Distance (mm) |
| --- | --- | --- | --- | --- |
| MiniVolt (this work) | 16.4 | 0.6 | 250 $\mu\text{m}$ ∅ | 1.5-1.6 |
| Miniscope-LFOV [27] | 13.9 | 0.25 | $3.6 \times 2.7$ mm | 3.5 |
| SimScope3D [23] | 6.65 | 0.3 | 207 $\mu\text{m}$ ∅ | 0.75 |
| UCLA miniscope (v4) [34] | 2.6 | N/A | 1 mm | 0.675-2 |
| MiniXL [24] | 3.5 | 0.12 | 3.5 mm | 1.9 |

MiniVolt recordings are conducted at 530 frames per second with an excitation intensity of 35-180 mW / mm^2^ using the Ximea CamTool GUI. Data is collected in Mono8 with pixels. Pixels are binned and the ROI is cropped to achieve voltage imaging framerates. We use 2 × 2 pixel binning, yielding an effective pixel pitch at the sample of 2.2 × 2.2 microns to enable full frame rate capabilities while still distinguishing cellular features.

### 2.2. Benchtop Imaging System

For benchtop measurements, following the diffusers described above, the 532 nm light is routed through a 1-meter multimode fiber patch cable (Thorlabs, FG600AEA). The light is imaged onto the sample with 90 mW / mm^2^ of intensity using a tube lens (Thorlabs, AC508-080-ML) and 16x/0.8NA long working distance (LWD) objective (Nikon, CFI75) and the emitted fluorescence is separated using a shortpass 567 nm dichroic mirror (Thorlabs, DMSP567L). Separated fluorescence passes through a long pass emission filter (Semrock, BLP01-568R-25) and 0.35× demagnification coupling (Olympus, U-TV0.35XC) before it is collected by a back-illuminated sCMOS (Hamamatsu Orca-Fusion BT, C15440-20UP) with an effective pixel pitch of 1.3 microns at the sample. The back aperture of the objective is restricted from M32 to M25, reducing the benchtop to ∼0.6 NA. This NA still enables sufficient PNR for voltage imaging, while creating a more realistic collection efficiency to compare with our miniature microscope. Once active cells have been identified on the benchtop system, the light is coupled into the MiniVolt multimode fiber to bypass the benchtop system, and mice are transitioned to head-fixation beneath the miniscope.

### 2.3. Sample Preparation

Transgenic NDNF-Cre mice (Jackson Laboratories, 028536 B6.Cg-Ndnftm1.1(folA/cre)Hze/J) were injected with 70 nl of the pGP-AAV-syn-FLEX-Volton2-ST-WPRE virus into the primary visual cortex (V1) (from Bregma: AP -3.71 mm and -3.5 mm, ML +2.5 mm and 2.0 mm, DV -0.15 mm) [15]. Subcutaneous injections of dexamethasone (0.25 mg/kg) and buprenorphine (0.1 mg/kg) in saline were used for pain management. After one month, a retro-orbital injection of Janelia Fluor 552 (JF552) was delivered and imaging was performed the following day. All animal experiments were approved by the University of Colorado Anschutz Medical Campus Institutional Animal Care and Use Committee (IACUC). Mice were first screened for expression using a benchtop widefield microscope (Fig. 2a) with a Hamamatsu ORCA Fusion-BT camera.

**Fig. 2.**
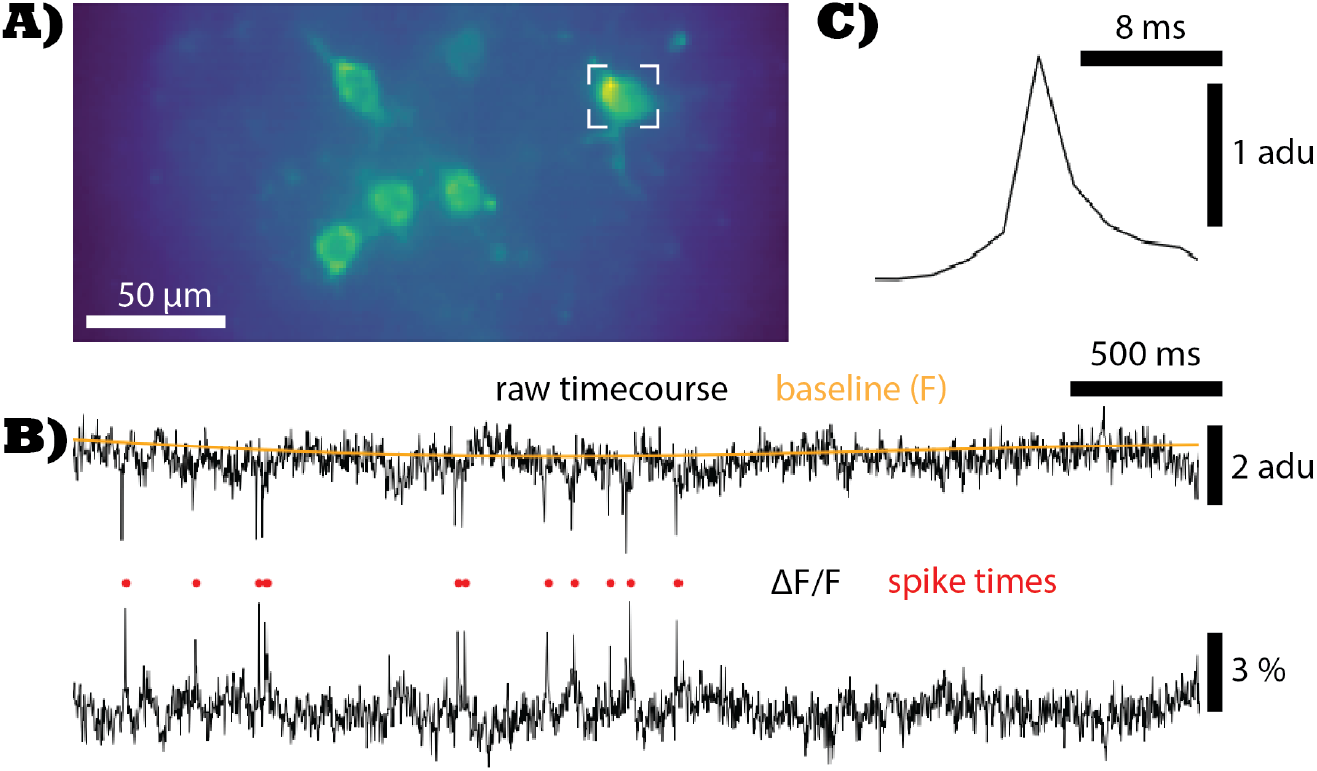
(a) Average intensity projection (AIP) of a 500 Hz MiniVolt recording of NDNF-Cre expressing cells in the visual cortex. (b) Raw intensity time course extracted using MiniVolt from the region of information marked in (a). Baseline fluorescence (orange) was calculated using a lowpass filter at 1/3 Hz to remove the effects of photobleaching without risking the removal of low frequency subthreshold oscillations in the theta band (4-12 Hz). ΔF / F time course (bottom) was obtained by dividing the raw signal by the baseline. Prominent spikes (indicated by red dots) are identified using the template in (c). (c) VolPy extracted spike template used to identify prominent spikes [6]

We selected the soma-tagged variant of Voltron2 (Voltron2-ST) to maximize our ability to distinguish dynamic background fluorescence from out-of-focus dendrites and in-focus cells. Unlike traditional calcium indicators, Voltron2 fluorescence decreases during spiking. This leads to a large constitutively fluorescent background that makes it challenging to measure the small fluorescense changes due to neural activity from in-focus cells and increases photon noise. By localizing emitted fluorescence to the soma, this out-of-focus background is decreased and easier to identify due to its lower spatial frequency.

### 2.4. Data Processing

Collected data either for benchtop or miniscope recordings (Fig. 2) are processed in the following steps. First, data are motion corrected using the VolPy implementation of non-rigid motion correction (NoRMCorre) [6, 35]. ROIs are segmented around in-focus cells in FIJI [36] (Fig. 2a) and extracted time courses (Fig. 2b) are detrended using a high pass filter at 1/3 Hz. A baseline is approximated by subtracting the high frequency information from the raw data (Fig. 2b, orange). ΔF / F time courses are obtained by dividing the raw data by the baseline. Resulting time courses are rectified (Fig. 2b), and spikes/spike templates (Fig. 2b-c) are identified using the VolPy implementation of SpikePursuit [6, 37]. Spike PNR is evaluated for each time course as transient ΔF/F height divided by the standard deviation during inactivity.

## 3. Results

Our goal is to demonstrate that MiniVolt is capable of voltage imaging in head-fixed mice and provide qualitative comparisons with a benchtop voltage microscope. An example voltage recording in a head-fixed mouse using MiniVolt is shown in Fig. 3. For the five labeled neurons, we provide ΔF / F time courses (Fig. 3b). A zoom in of the time course for neuron 3, the most active neuron in this window, is shown with corresponding spike times (Fig. 3c). The spike ΔF / F is around 2-5% and the PNR is 4.2, which is on par with benchtop widefield imaging. We characterized the minimum excitation to observe spiking in MiniVolt with a PNR threshold of 3 is 35 mW / mm^2^ (see Fig. S2).

**Fig. 3.**
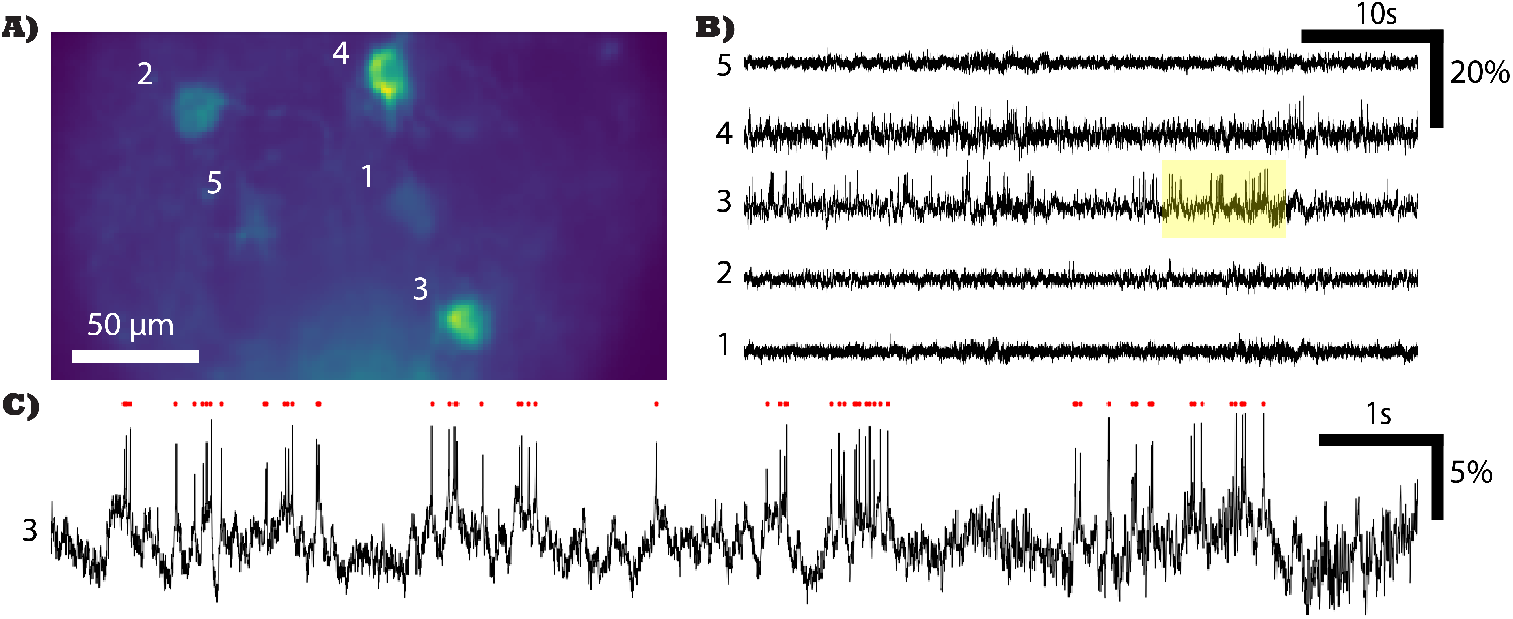
Example voltage recording using MiniVolt in a head-fixed mouse. (a) AIP of NDNF interneurons expressing Voltron2_552_ in the visual cortex. Data was recorded at 530 Hz with 170 / mW mm2 intensity at the sample. (b) ΔF / F time courses for 5 regions of information indicated in (a). (c) Zoomed-in time course for the yellow highlighted region in (b). The red points above the time course are the timing of spiking events identified using VolPy

A comparison of the performance between MiniVolt and our benchtop widefield microscope is shown in Figure 4. Figures 4a and 4b show an average intensity projection (AIP) and extracted ΔF / F time courses for a benchtop widefield recording of Voltron2 expressed in NDNF interneurons the mouse visual cortex. Figures 4c and d are the AIP and ΔF / F time courses from MiniVolt in the visual cortex. These results demonstrate the comparable PNR in recordings taken using the benchtop and MiniVolt (Fig 4e). VolPy extracted spike templates for benchtop and MiniVolt taken from ROI 1 in each recording are shown for comparison in Fig. 4f. It is important to note that different excitation intensities were used (90 mW / mm^2^ for the benchtop and 180 mW / mm^2^ for MiniVolt) and PNR increases with intensity [15]. Additionally, different cells were imaged in the two systems so the comparison is qualitative.

**Fig. 4.**
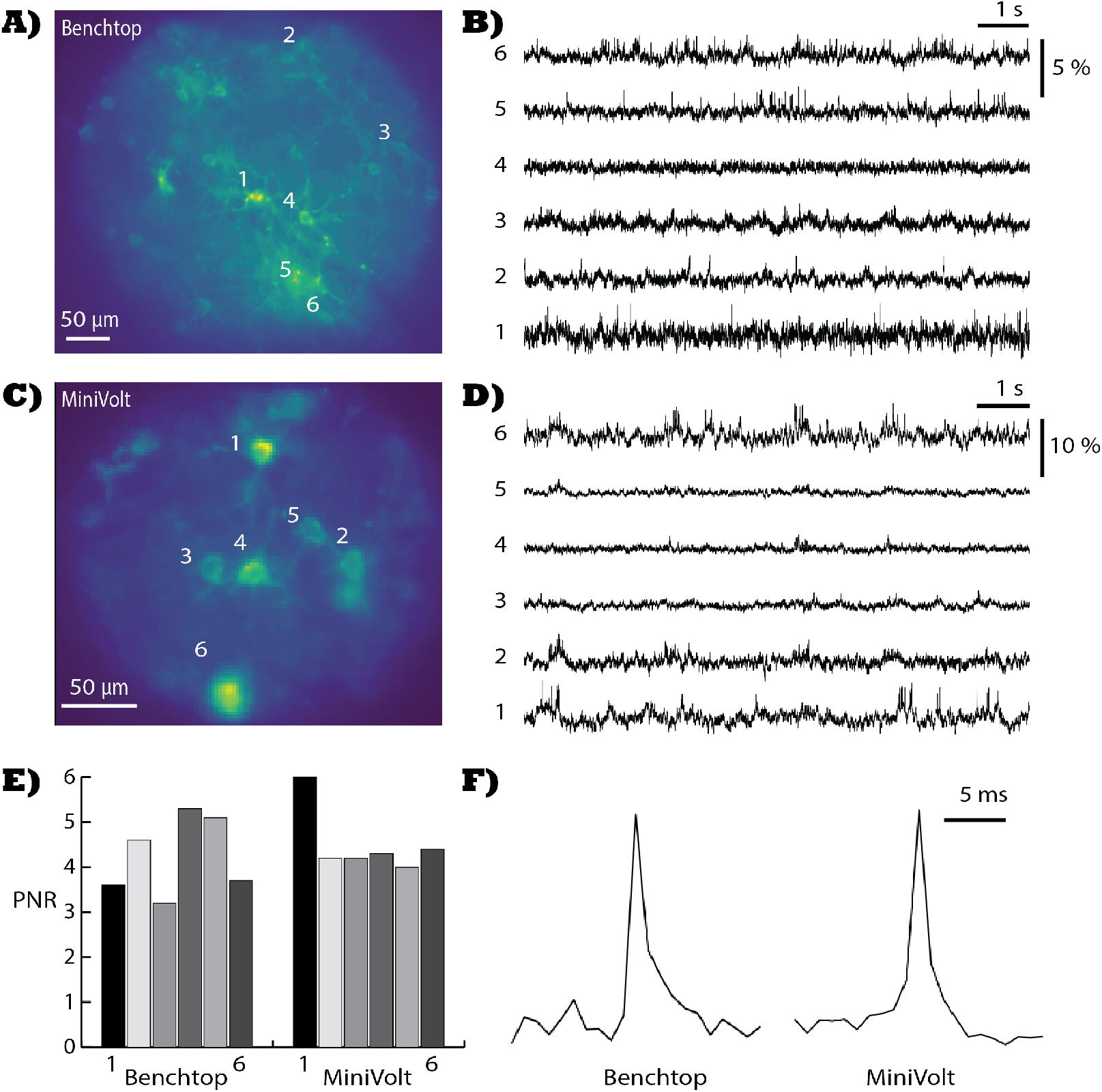
Example benchtop and MiniVolt voltage recordings. (a) AIP of a benchtop recording taken at 500 frames per second with 90 mW / mm^2^intensity. (b) ΔF /F time courses for 6 ROIs indicated in (a). (c) AIP of a MiniVolt recording taken at 525 frames per second with 180 mW / mm^2^intensity. (d) ΔF /F time courses for 6 ROIs indicated in (c). (e) Mean PNR for the time courses for ROIs shown in (b) and (d). (f) VolPy spike templates extracted from the benchtop recording (ROI1) and the MiniVolt recording (ROI1).

## 4. Discussion

Collection efficiency is an important microscope design constraint for voltage imaging. Voltage indicators rely on detecting small (<10%) changes in baseline fluorescence signal (this can be less than 2% change when contaminated with out-of-focus background), and thus, a sufficient number of photons must be collected in each frame in order to resolve these small changes over the background and noise. For widefield imaging, the baseline signal will increase linearly with excitation intensity. Based on the frame rate of the system, we characterized the minimum intensity needed to distinguish changes in in-focus signal from shot noise to be ∼ 35 mW / mm^2^ for a frame rate of 531 Hz using MiniVolt (Fig. S2). However, phototoxicity, photodamage, and photobleaching place an upper bound on the usable intensity, which we experimentally found to be (∼ 200 mW / mm^2^). It is therefore important to maximize the collection efficiency of MiniVolt to operate below this threshold. We were able to aperture down our benchtop to obtain a 0.6 NA objective while still preserving a high PNR in our voltage imaging. The filter set efficiency is less for the benchtop (63%) compared to MiniVolt (71%), which helped with collection efficiency, even with the reduced numerical aperture of 0.6 in MiniVolt. In order to achieve a higher NA in our MiniVolt system compared with the UCLA Miniscope V4, it was necessary to use larger diameter lenses and a water immersion design.

We selected a miniature camera with a SONY IMX568 image sensor due to its pixel size (2.74 × 2.74 µm), quantum efficiency (> 60% for 550-650 nm) and back-illuminated and stacked CMOS structure. Harnessing Pregius S technology, the IMX568 is optimized for high sensitivity and low-noise recording despite small pixel sizes. Smaller pixel sizes were required because the magnification of an imaging system depends on the difference in focal lengths of the objective and tube lenses, which is why miniature microscope magnification is limited, typically ∼ 1.5-2× due to size constraints. The industry standard Miniscope V4 utilizes an onboard Python480 image sensor. Compared to the Sony IMX568, the Python480 has 24% less quantum efficiency at 550 nm, decreased framerate capabilities, larger pixels (4.8 × 4.8 µm) and decreased dynamic range. Table S1 shows the quantum efficiency, dynamic range, full well capacity, pixel size and read noise for these two image sensors along with the Aptina MT9P031 (SIMScope3D and MiniXL) and the high-end camera used on our benchtop microscope for voltage imaging (Hamamatsu Orca Fusion-BT).

MiniVolt is the only miniaturized widefield microscope with an onboard camera capable of voltage imaging at framerates > 1000 Hz. Due to limitations in excitation intensity and PNR, the maximum framerate we demonstrate in this work is 530 Hz. Although our device has not been demonstrated on freely moving animal models, the size and weight is comparable with miniscopes that have been used in freely moving imaging in rat models, such the large field of view Miniscope-LFOV [27]. In comparison with other miniscopes, MiniVolt has the highest collection efficiency and sensor speed which enables voltage imaging.

To extend voltage imaging to freely moving imaging in all rodent models, it will be necessary to further reduce the weight in our design. Our characterization has revealed that high numerical aperture is crucial for high fidelity voltage imaging. Thus, custom optics will be necessary to enable smaller diameter lenses while preserving the numerical aperture. This has been a successful strategy in two- and three-photon miniscopes [29, 38]. Additionally, sensor technology is constantly progressing. The camera used herein was released only a year ago. In MiniVolt, we are using a small fraction of the overall sensor. Development of a CMOS sensor with similar specifications but smaller size would significantly help to reduce weight along with custom small format CMOS image sensor printed circuit boards (PCB).

## 5. Conclusion

MiniVolt represents a significant advancement in miniaturized widefield microscopy, specifically engineered for high-speed voltage imaging in freely moving animals. This innovative device addresses the critical challenges associated with observing neural activity, particularly the necessity for high collection efficiency and rapid image sensors to accurately capture the subtle changes in fluorescence produced by voltage spikes. We demonstrate a 0.6 numerical aperture miniscope (MiniVolt) that incorporates a high speed camera (capable of >1 kHz frame rate) with photon collection efficiency to enable in vivo voltage recording of neural activity at 530 Hz. MiniVolt is validated by performing voltage recording from head-fixed NDNF-Cre transgenic mice injected with Voltron2. Recordings acquired on MiniVolt and a standard widefield benchtop microscope show comparable voltage time courses from individual NDNF interneurons with similar peak to noise ratios. MiniVolt uses only off the shelf commercial optics that have a lower cost than custom designs to achieve high numerical aperture and additionally combines a commercial low form factor CMOS camera with high quantum efficiency and framerate. With a total device weight of 16.4 grams, MiniVolt is comparable in weight to miniscopes developed for imaging in freely moving rats. Future work can reduce the weight further by including custom optics to maintain the high numerical aperture with smaller diameters. Additionally, with the progress in sensor technology, use of a smaller size sensor combined with small format PCB can enable further miniaturization to extend freely moving voltage imaging in all rodent models.

## Data availability

Data underlying the results presented in this paper are not publicly available at this time but may be obtained from the authors upon reasonable request.

## Supplemental document

See Supplement 1 for supporting content.

