## Supplemental Materials for "Miniaturized widefield microscope for high speed in vivo voltage imaging"

**Table S1.**

| Image Sensor | QE @ 550nm (%) | Dynamic Range (dB)* | Full Well Capacity (e-) | Pixel Size (micron) | Read Noise (e-) |
|---|---|---|---|---|---|
| IMX568 | 80% | 71 | 9,200 | 2.74 | 2.6 |
| Python480 (Miniscope V4) | 56% | 59 | 10,000 | 4.8 | 11.0 |
| Fusion BT (fast scan) | 96% | 79 | 15,000 | 6.5 | 1.6 |
| MT9P031 (SIMScope3D and MiniXL) | 60% | 60 | 4,900 | 2.2 | 5.0 |

*calculated from ratio between Full Well Capacity and Read Noise

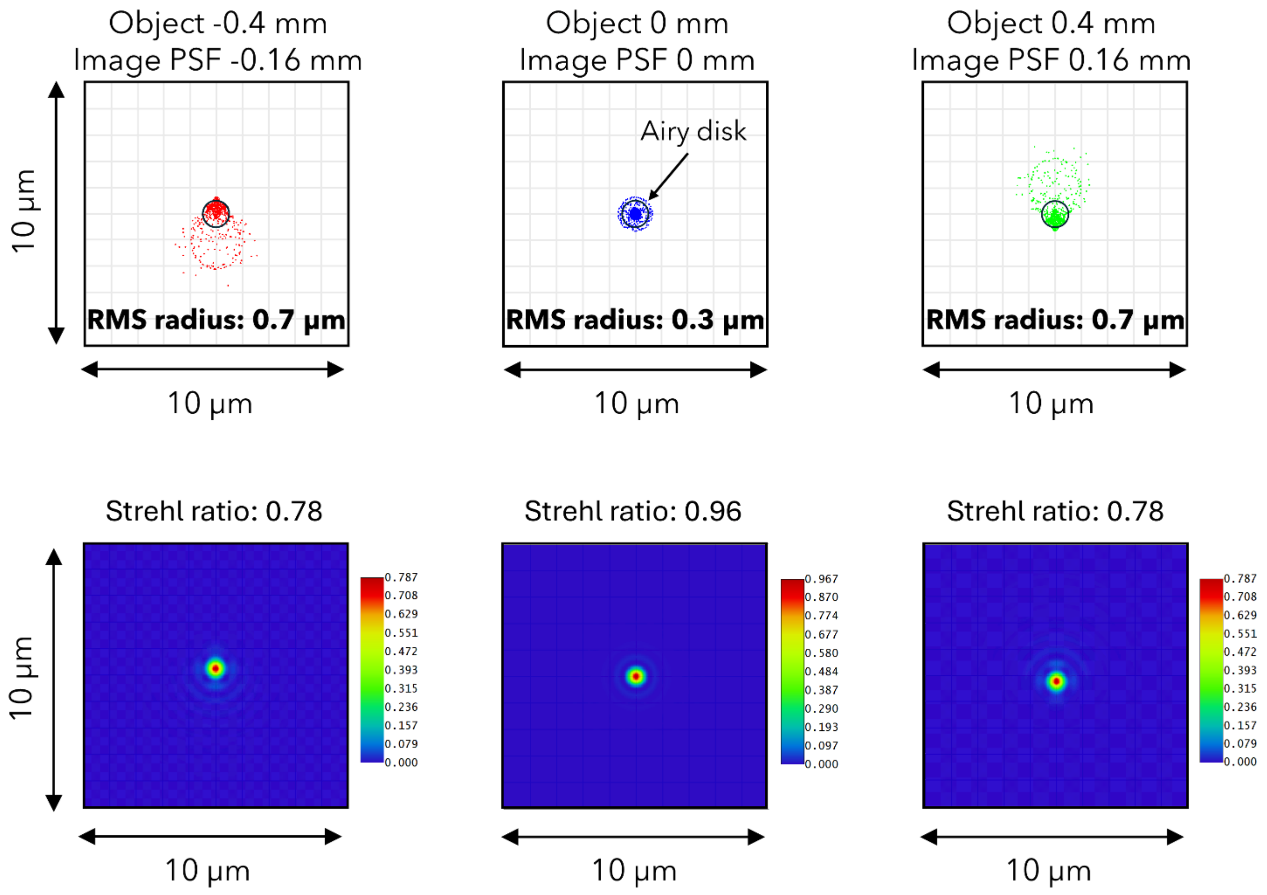


**Figure S1. Simulations of the MiniVolt point-spread function.** The spot size and point-spread function (PSF) are shown for an object on- and off-axis for MiniVolt.

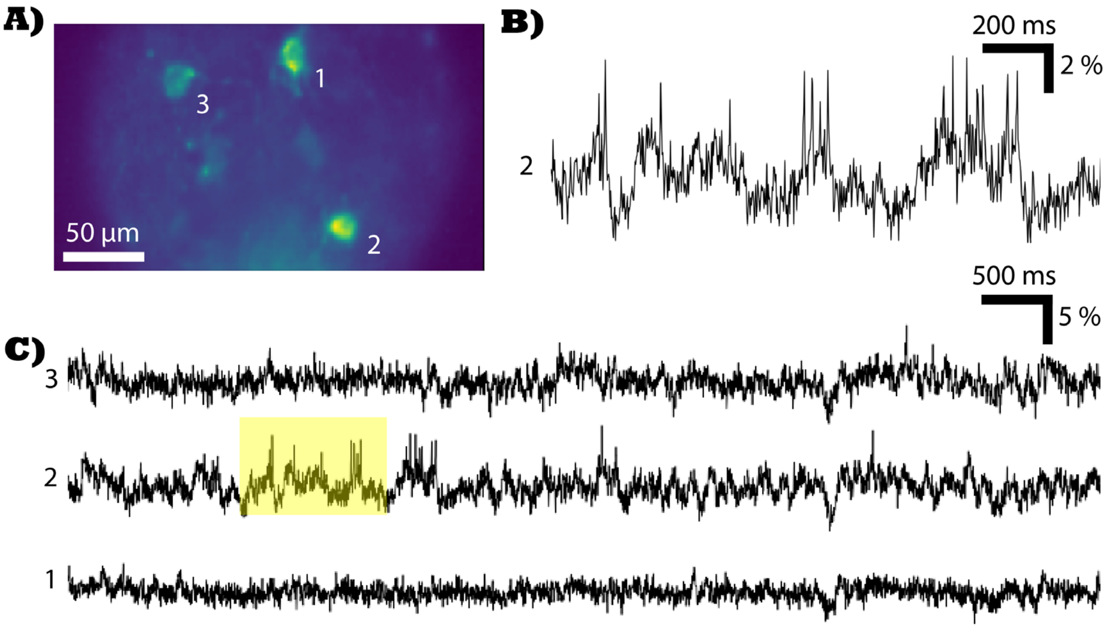


**Figure S2. Voltage recording from MiniVolt at low excitation power.** (a) Average intensity projection of NDNF interneurons expressing Voltron2552 in the visual cortex of an awake mouse. Data was recorded at 531 Hz with 35 mW/mm$^2$ intensity at the sample. (b) ΔF/F time course from ROI 2 indicated in (a). (c). ΔF/F time course from three different ROIs indicated in (a).
